# Gene Expression Mediated Antiproliferative Potential and Safety of Selected Medicinal Plants Against Cancerous and Normal Cells

**DOI:** 10.1101/578948

**Authors:** Regina Wachuka Mbugua, Eliud Mwaniki Njagi, Chrispus Mutuku Ngule, Peter Mwitari

## Abstract

Globally, approximately 13% of all deaths annually are attributed to cancer. Surgery, radiation and chemotherapy are the current treatment techniques for cancer, however these methods are expensive, have high failure rates and have been associated with detrimental side effects. Plant derived products could be good candidates in alleviating challenges being experienced with these current methods. This study aimed at evaluating the phytochemistry, antiproliferation potential, and probable mechanism of action of *Albizia gummifera*, *Rhamnus staddo* and *Senna didymobotrya* plant extracts. Phytochemical screening was done as per standard procedures. The common 3– (4, 5-dimethylthiazol-2-yl) -2, 5-diphenyltetrazolium (MTT) dye was used in the determination of the antiproliferative activity of the extracts. Extracts induction of *VEGF* (angiogenesis) and p53 (apoptosis) genes’ expression was evaluated using Real Time Polymerase Chain Reaction. Phytochemical screening revealed presence of alkaloids, tannins, glycosides, flavonoids, terpenes, phenolics and saponins in the plants extracts. *A. gummifera’s* stem bark methanol: dichloromethane extract had the highest activity against the cancerous cell lines tested: HCC1395 (IC_50_ 6.07±0.04μg/ml), DU145 (IC_50_ 3.34±0.05μg/ml), CT26 (IC_50_ 5.78±0.08μg/ml) and Hep2 (IC_50_ 7.02±0.01μg/ml). *R. staddo* root bark methanol: dichloromethane extract had an IC_50_ value of 15.71±0.04μg/ml on HCC, 9.81±0.09μg/ml on Hep2 and 11.14±0.39μg/ml on CT26. *S. didymobotrya* root bark methanol: dichloromethane extract inhibited HCC with an IC_50_ of 65.06±0.07μg/ml, CT26 with an IC_50_ of 15.71±0.04μg/ml and Hep2 with an IC_50_ of 62.10±0.11μg/ml. From the results obtained, the plants exhibited selective toxicity to cancer cells while sparing the normal cells (SI ≥ 3). *A. gummifera* and *S. didymobotrya* and *R. staddo* plant extracts upregulated p53 and down-regulated *VEGF* genes. In conclusion, this study confirms that these plant extracts could be potential candidates for development of drugs for the management of breast, prostrate, colorectal and throat cancer.

## INTRODUCTION

Cancer is among the leading causes of morbidity and mortality worldwide. Approximately 14 million new cases and 8.2 million cancer related deaths occurred in 2012 [1]. Cancer new cases are anticipated to rise by 70% in the next two decades [1]. In Kenya, cancer ranks third after cardiovascular and infectious diseases as cause of mortality. Approximately, 28,000 cases are recorded annually [2]. The factors that have been associated with high cancer risk include high body mass index, low fruit and vegetable intake, lack of physical activities, environmental pollution, tobacco use and alcohol [1].

Chemotherapy, radiation, surgery and hormonal therapy are the main strategies employed in the management of cancer. Despite their effectiveness, they have various side effects such as hair loss, peripheral neuropathy and cardiac damage among others [3]. Due to these challenges, people have turned to the use of medicinal plants as alternative therapies because they are thought to be cheap, effective, safe and easily accessible [3]. It has been estimated that over 30% of the plant’s species contain secondary metabolites which are useful in treatment of various diseases such as cancer [3]. The use of naturally derived products from medicinal plants that selectively induce apoptosis and reduce angiogenesis could serve as an alternative to the current cancer treatment regimes. The discovery of cancer treatment drugs vincristine and vinblastine from *Catharanthus roseus* has prompted medicinal plants research for new leads in cancer treatment and management [4].

*S. didymobotrya* belongs to the family Fabaceae (Leguminosae) [5]. An ethnobotanical study conducted in Kakamega County in Kenya, showed that the leaves of *S. didymobotrya* are boiled and taken together with the leaves of *Galinsoga parviflora*, *Triumfetta rhomboidea* and *Ocimum gratissimum* for the treatment of colorectal cancer [6]. Elsewhere in Kenya, the plant has been used traditionally by the Kipsigis community in the control of malaria as well as diarrhea [7]. In West Pokot, the pastoralists peel the bark, dry the stem and burn it into charcoal which they use in milk preservation [5]. The root decoction has been used in several countries such as Congo, Rwanda, Burundi, Kenya, Tanzania, and Uganda to treat malaria, ringworms, jaundice and intestinal worms [8].

*A. gummifera* belongs to the Leguminosae (Fabaceae) family and subfamily Mimosoideae [9]. The Albizia species have numerous ethnomedical uses. *A. gummifera* has been widely used traditionally in African countries [10]. Traditionally, the leaf and stem bark are boiled together with the leaf of *Salivia coccinea* and *Conyza sumatrensis* and taken orally for the treatment of colorectal cancer, throat cancer, breast cancer and squamous cell carcinoma of the gums [6]. The bark infusion and pounded bark is taken to treat malaria and snuffed to treat headache, respectively, in Kenya. An aqueous acetone extraction on *A. gummifera* showed *in vitro* anthelminthic activities of condensed tannins on egg hatchability and larval development of sheep *Haemonchus contortus* [11]. Compounds isolated from *A. gummifera* exhibited cytotoxicity against the A2780 human ovarian cancer cell lines [12].

*R. staddo* belongs to family Rhamnaceae. It is commonly known as staddo or buckthorn. It has been shown that plants from the family Rhamnaceae possess anticancer activity [13]. *Ziziphus spina christi* is an example of a plant belonging to this family. It demonstrated anti-proliferative potential on MCF-7 breast cancer cells by inducing apoptosis [14]. Traditionally, its roots are used to treat typhoid, back pain, joint pain and headache. The leaf is burned and smelt to increase male libido [15]. The root bark is also used by the Kenyan Maasai community to extend the shelf life of milk [16]. *R. staddo* has been reported to have both antifungal and antimalarial activity when used synergistically with other plants [17].

Several important bioactive compounds that produce desirable physiological activities have been derived from plants. These plants could serve as new leads and clues for modern drug design [18]. These important bioactive constituents of plants include but are not limited to alkaloids, tannins, flavonoids, terpenoids and phenolic compounds [19]. During the synthesis of compounds with specific activities to treat various diseases such as cancer, it is important to know the correlation between the phytoconstituents and the bioactivity of plants [20]. Therefore, preliminary screening of plants is the need of the hour in the discovery and development of safe novel therapeutic agents with improved efficacy.

The determination of the molecular mechanisms underlying neoplastic transformation and progression have resulted in the understanding of cancer as a genetic disease, which evolved from the accumulation of a series of acquired genetic lesions [21]. Protein 53 (p53) is a tumor suppressor that eliminates and inhibits multiplication of abnormal cells through induction of apoptosis [22]. It is one of the key orchestrators of cell signaling pathways related to apoptosis and cell cycle, which have an essential role in the development and progression of complex diseases such as cancer [23]. Studies have shown that medicinal plants can activate apoptotic genes [24].

Angiogenesis is a key process in cancer promotion. It is an important pathological event associated with tumor growth and metastasis. Vascular Endothelial Growth Factor (*VEGF*) plays an important role in this event [25]. It is a physiological process of formation of new blood vessels on already existing ones. The newly formed blood vessels facilitate the metastatic dissemination of cancer cells. In most cancers, angiogenesis correlates with disease stage and metastasis [26].

Recently, the role of Vascular Endothelial Growth Factor (VEGF) in angiogenesis has attracted the attention of researchers. This gene plays an important role in regulation of physiological as well as pathological angiogenesis [27]. Various reports have shown that plant extracts and plant derived compounds have the potential to down regulate VEGF. Octacosanol, a long-chain aliphatic alcohol derived from *Tinospora cordifolia* suppressed the expression of VEGF an activity which was attributed to the compounds ability to regulate IL-1β, IL-6 and TNF-α cytokines levels [28, 29]. *Withania somnifera* inhibits VEGF mediated neovascularization [30] and Z-guggulsteron isolated from *Commimpora mukul* plant down regulated angiogenesis through VEGF and granulocyte colony-stimulating factor, pro-angiogenic growth factors inhibition [31]. In most cancers, angiogenesis correlates with disease stage and metastasis [32]. Metastasis is the major cause of death from cancer. The development of rich vascular networks of new lymphatic and blood vessels in the tumors results in resistance to conventional therapies such as androgen ablation and cytotoxic chemotherapy [33].

## Materials and Methods

### Study site

The study was carried out at the Center for Traditional Medicine and Drug Research (CTMDR), Kenya Medical Research Institute (KEMRI) Nairobi, Kenya.

### Study design

*In vitro* laboratory based (pre-clinical) and *in vivo* experimental study design method was used.

### Sample collection and preparation

The leaf, stem bark and root bark of *Senna didymobotrya* and *Rhamnus staddo* were collected from Laikipia County. *Albizia gummifera* plant parts were collected from Ngong Forest, Kajiado County. Harvesting was done sustainably. Identification of the botanical samples was conducted by a qualified botanist and voucher specimens (RAM 2017/01, RAM 2017/03 and RAM 2017/2 respectively) stored at the University of Nairobi Herbarium. The leaf, stem bark and root bark collected were dried at room temperature and then ground into fine powder using Gibbons electric mill (Wood Rolfe Road Tolles Bury Essex, UK). The ground samples were then stored in air tight bags at room temperature until use.

## Extraction

### Aqueous Extraction

About 200g of each sample was weighed and submerged in 1litre of double distilled water. Extraction was done in an aqueous bath at 80^0^C for 2hrs. After cooling, the extract was decanted in a clean 1000ml conical flask and filtered using a Whatman No. 1 filter paper. The filtrate was then freeze dried using a freeze dryer (Modulyo Edwards high vacuum, Crawey England, Britain, Serial No. 2261). The extract was weighed and stored at 4^°^C in air tight vials until use [34].

### Organic Extraction

Briefly, 200g of each sample was weighed, put in a flat-bottomed conical flask and solvent added to cover the sample completely and left to stand for 24hrs. A Whatman No. 1 filter paper was used to filter and the sample re-soaked again for 24hrs. Extraction was done using methanol: dichloromethane (1:1). The solvents were removed using a rotary evaporator (Büchi, Switzerland) and the concentrated extracts packed in air tight vials and stored at 4^°^C until use [27].

### Qualitative Phytochemical Screening

Qualitative phytochemical screening of *S. didymobotrya*, *A. gummifera* and *R. staddo* was done using standard procedures [35]. Secondary metabolites tested included alkaloids, tannins, steroids, glycosides, flavonoids, phenols, saponins, and terpenoids.

### Cell culturing

DU 145 (prostate cancer), HCC 1395 (breast cancer) Hep 2(throat cancer), CT26 (colorectal cancer) and Vero E6 (normal) cells obtained from ATCC (Manassas, VA, USA) were used. The cells initially stored in liquid nitrogen were removed from the tank and quickly thawed in a water bath at 37°C. The vial contents were centrifuged, supernatant removed and then transferred into growth MEM medium supplemented with 10% Fetal Bovine Serum (FBS), 1% L-Glutamine and 1% antibiotic (Penicillin/Streptomycin) in a T75 culture flask and incubated at 37°C and 5% CO_2_ to attain confluence.

### Antiproliferative assay

Upon attainment of confluence, cells were washed with saline phosphate buffer and harvested by trypsinization. The number of viable cells was determined using Trypan blue exclusion method (cell density counting) using a hemocytometer. An aliquot of 100μl containing 2.0 ×10^4^ cells/ml suspension was seeded in to a 96-well plate and incubated at 37^°^C for 24hrs at 5% CO_2_ for 24hrs. After 24hrs, 15μl of sample extracts at seven different concentrations each serially diluted were added on Row H-B. Row A, containing media and cells alone served as the negative control. The standard drug Doxorubicin was used as the positive control. The experiment was done in triplicate. The cells were incubated for 48hrs, then 10 μl of MTT dye (5mg/ml) was added and the plates incubated for 2hrs at 37°C and 5% CO_2_. Mitochondrial dehydrogenase which is a biomarker of live cells interacts with MTT dye reducing it to insoluble formazan. The formazan formed is directly proportional to the number of live cells. Formazan formation was confirmed using inverted light microscope and then solubilized with 50μl of 100% DMSO and optical density (OD) read using a calorimetric reader at 540nm and a reference wavelength of 720nm. The effect of the test samples on the cancer and normal cells was expressed as IC_50_ values (the extracts concentration which kills 50% of the cancer cells) and CC_50_ values (concentration of extracts that exerted cytotoxic effects to 50% of the normal cells) respectively [36].

Selectivity index (SI) which indicates the ability of the extracts to exert selective toxicity to cancer cells while sparing the normal ones was calculated using the formula:

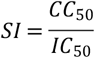

Where;

CC_50_ – Concentration of extract that exerted cytotoxic effect to 50% of the normal cells

IC_50_– Concentration of extract that inhibited the growth of cancer cells by 50%

The data obtained was analysed using linear regression model to get IC_50_ of each drug. The IC_50_ values of the extracts were compared using Minitab Version 18 to obtain the Mean±SEM.

### RNA Extraction and Gene Expression Assay Cell Culture and RNA Extraction

The prostate cancer cell line (DU 145) was used in the evaluation of the gene expression profiles of the most active plant extracts from each of the three plants. The cell line used was selected based on the antiproliferative effect of the extracts. The cells obtained from ATCC (Manassas, VA, USA) were cultured in a T-25 culture flask. Minimum Essential media (MEM) supplemented with 10% Fetal Bovine Serum (FBS), 1% L-Glutamine and 1% antibiotic (Penicillin/Streptomycin) was then added and the flask incubated at 37°C and 5% CO_2_ to attain confluence. After 24hrs, *S. didymobotrya*, *A. gummifera* and *R. staddo* extracts were added to the T-25 flask at appropriate concentrations. The concentrations of the extracts used in this assay were informed by the IC_50_ values earlier obtained. To get the concentration of the extract to expose to the cultured cells and get enough viable cells, the IC_50_ values were reduced by 30%. This was then incubated for 48 hours. After 48hrs, the media was decanted and cells washed in PBS to remove any debris. Trypsinization of the cells was done. RNA extraction was carried out using the procedure described by Pure Link RNA mini kit (Thermo Scientific, USA). The extracted RNA was quantified and its quality examined using a Nanodrop ND-2000 spectrophotometer (Nanodrop Technologies, Inc., Wilmington, DE, USA) and the concentrations (ng/μl) obtained.

### Gene Expression Assay

Reverse transcription and cDNA amplification were done in single step reaction using SuperScript IV Reverse Transcriptase and Thermo scientific Real time SYBR green Kits according to the manufacturer’s instructions. A single narrow peak from each PCR product was obtained by melting curve analysis at specific temperatures. The quantitative RT-PCR data was analyzed by a comparative threshold (Ct) method, and the fold inductions of the samples compared with the untreated samples. Glyceraldehyde 3-phosphate dehydrogenase was used as an endogenous control to normalize the expression of the target genes. The Ct cycle was used to determine the expression level in control and cells treated with different extracts for 48hours. The geneexpression levels were calculated as described by Yuan *et al* [30]. The data obtained was expressed as the ratio of the reference gene to the target gene using the standard formula: ΔCt = Ct (test gene) − Ct (GAPDH). To determine the relative expression levels, the following formula was used: ΔΔCt = ΔCt (treated) −ΔCt (calibrator). Therefore, the expression levels were presented as *n*-fold differences relative to the calibrator. The value was used to plot the expression of apoptotic and angiogenic genes using the relative quantification (2^-ΔΔCt^) [37].

## Results

### IC_50_ and CC_50_ values for the plants extracts

On the prostate cancer cell line (DU 145), the stem bark extract of *A. gummifera* MeOH: DCM exhibited the highest cell inhibitory effects with an IC_50_ value of 3.34±0.05μg/ml followed by *R. staddo* root bark methanol dichloromethane and *A. gummifera* aqueous stem bark extracts at IC_50_ values of 9.36±0.10μg/ml and 18.29±0.02μg/ml, respectively (Table 1). *A. gummifera* MeOH: DCM root bark had the lowest activity exhibiting the lowest IC_50_ value of 79.71±0.10μg/ml. Amongst all the *S. didymobotrya* extracts tested on the prostate cancer cell line, only the leaf MeOH: DCM extract portrayed activity with an IC_50_ value of 65.72±0.01μg/ml (Table 1). There was a significant difference among the IC_50_ values of all the extracts and the positive control (p≤0.05) against DU 145 cell line.

**Table 1:** IC_50_and CC_50_values of the plant extracts on the selected cell lines

| Plant Sample | Part Used | Solvent | DU145<br>IC<sub>50</sub> (µg/ml) | HCC 1395<br>IC<sub>50</sub> (µg/ml) | Hep 2<br>IC<sub>50</sub> (µg/ml) | CT 26<br>IC<sub>50</sub> (µg/ml) | VERO<br>CC<sub>50</sub> (µg/ml) |
| --- | --- | --- | --- | --- | --- | --- | --- |
| *S. didymobotrya* | Stem bark | Aqueous | >100 | >100 | >100 | >100 | >100 |
|  | Leaves | Aqueous | >100 | >100 | >100 | >100 | >100 |
|  | Root bark | Aqueous | >100 | >100 | >100 | >100 | >100 |
|  | Stem bark | MeOH: DCM | >100 | 58.67±0.02<sup>b</sup> | 66.66±0.35<sup>a</sup> | >100 | >100 |
|  | Leaves | MeOH: DCM | 65.72±0.01<sup>c</sup> | 32.32± 0.03<sup>e</sup> | 39.17±0.29<sup>c</sup> | >100 | >100 |
|  | Root bark | MeOH: DCM | >100 | 65.06±0.07<sup>a</sup> | 62.10±0.11<sup>b</sup> | 15.13±0.03<sup>d</sup> | >100 |
| *A. gummifera* | Stem bark | Aqueous | 18.29±0.02<sup>f</sup> | 21.38±0.03<sup>f</sup> | 23.41±0.21<sup>f</sup> | 27.00±0.12<sup>c</sup> | 63.00±0.0<sup>d</sup> |
|  | Leaves | Aqueous | 66.26±0.04<sup>b</sup> | >100 | >100 | >100 | >100 |
|  | Root bark | Aqueous | 25.29±0.09<sup>e</sup> | 35.58±0.25<sup>d</sup> | 55.38±0.38<sup>c</sup> | 64.77±0.13<sup>b</sup> | 84.43±0.4<sup>b</sup> |
|  | Stem bark | MeOH: DCM | 3.34±0.05<sup>h</sup> | 6.07±0.04<sup>h</sup> | 7.02±0.01<sup>i</sup> | 5.78±0.08<sup>g</sup> | 15.68±0.0<sup>f</sup> |
|  | Leaves | MeOH: DCM | 64.48±0.24<sup>d</sup> | 53.77±0.06<sup>c</sup> | 49.21±0.33<sup>d</sup> | 79.86±0.14<sup>a</sup> | 85.64±0.0<sup>e</sup> |
|  | Root bark | MeOH: DCM | 79.71±0.10<sup>a</sup> | 2.38±0.019<sup>i</sup> | 12.15±0.07<sup>g</sup> | 6.04±0.17<sup>f</sup> | 52.27±0.3<sup>e</sup> |
| *R. staddo* | Root bark | MeOH: DCM | 9.36±0.10<sup>g</sup> | 15.71±0.04<sup>g</sup> | 9.81±0.09<sup>h</sup> | 11.14±0.39<sup>e</sup> | 77.95±0.1<sup>c</sup> |
| **Doxorubicin** |  |  | 0.24±0.03<sup>i</sup> | 0.54±0.30<sup>j</sup> | 0.26±0.01<sup>j</sup> | 2.95±0.04<sup>h</sup> | 0.98±0.01<sup>g</sup> |

Values are expressed as Mean±SEM. Values that do not share a letter are significantly different (p≤0.05).

On the breast cancer cell line (HCC 1396), *A. gummifera* root bark MeOH: DCM extract had the highest cell inhibition with an IC_50_ value of 2.38±0 .01μg/ml. This was then followed by the *A. gummifera* stem bark MeOH: DCM extract *R. staddo* root bark MeOH: DCM extract with IC_50_ values of 6.07±0.04μg/ml and 15.71±0.04μg/ml respectively. Among the *S. didymobotrya* extracts, the leaf MeOH: DCM extracts had the greatest activity with an IC_50_ value of 32.32± 0.03μg/ml. *A. gummifera* stem bark and root bark aqueous extracts exhibited cell growth inhibition with IC_50_ values of 21.38±0.03μg/ml and 35.58±0.25μg/ml, respectively (Table 1). Amongst all the extracts tested on the HCC 1396 cell line, *S. didymobotrya* root bark MeOH: DCM extract had the lowest activity with an IC_50_ value of 65.06±0.07μg/ml (Table 1). A significant difference was observed between all the extracts and the positive control (p≤0.05).

On the throat (larynx) cancer cell line (Hep 2), the *A. gummifera* extracts, the MeOH: DCM stem bark and root bark extracts were shown to have IC_50_ values of 7.02±0.01μg/ml and 12.15±0.07μg/ml, respectively. *A. gummifera* stem bark aqueous extract had an IC_50_ value of 23.41±0.21μg/ml. The root bark MeOH: DCM extract of R. staddo also inhibited the growth of the throat cancer cells with an IC_50_ value of 9.81±0.09μg/ml. *S. didymobotrya* leaf MeOH: DCM extract exhibited an IC_50_ value of 39.17±0.29μg/ml. The stem bark MeOH: DCM extract of *S. didymobotrya* showed the lowest activity with an IC_50_ value of 66.66±0.35μg/ml (Table 1). There was significant difference among the IC_50_ values of all the extracts and the positive control (Table 1; p≤0.05).

*A. gummifera* stem bark and root bark MeOH: DCM extracts had the highest inhibition on colorectal (CT26) cancer cell line with IC_50_ values of 5.78±0.0μg/ml and 6.04±0.17μg/ml respectively. *A. gummifera* stem bark aqueous extract had an IC_50_ value of 27.00±0.12μg/ml. Root bark MeOH: DCM extract of *R. staddo* also displayed an IC_50_ value of 11.14±0.39μg/ml (Table 1). The root bark MeOH: DCM extract of *S. didymobotrya* showed the highest an IC_50_ value of 15.13±0.03μg/ml against colorectal (CT26) cancer cell line. The lowest cell inhibition was shown by *A. gummifera* leaf MeOH: DCM extract at an IC_50_ value of 79.86±0.14μg/ml. A significant difference was observed among the IC_50_ values of all the extracts and the positive control (p≤0.05). The CC_50_ values of the plant extracts varied depending on the plant extract and the solvent of extraction used. It was observed that most of the aqueous extracts were not cytotoxic (CC_50_≥100) and compared to the MeOH: DCM plant extracts. All the *S. didymobotrya* extracts and the *A. gummifera* leaf aqueous extract had CC_50_ values greater than 100μg/ml. Stem bark MeOH: DCM extract of *A. gummifera* was the most cytotoxic with a CC_50_ value of 15.68±0.08μg/ml. This was then followed by the root bark MeOH: DCM and the stem bark aqueous extracts of the same plant with CC_50_ values of 52.27±0.37μg/ml and of 63.00±0.01μg/ml. *R. staddo* root bark MeOH: DCM extract displayed a CC_50_ value of 77.95±0.14μg/ml. The lowest cytotoxicity was observed in leaf MeOH: DCM and root aqueous extracts of *A. gummifera* with CC_50_ values of 85.64±0.07μg/ml and 84.43±0.49μg/ml respectively. The doxorubicin was the most toxic towards the normal cells with a CC_50_ value of 0.98±0.01μg/ml (Table 1). All Aqueous and MeOH: DCM leaf extracts of *S. didymobotrya* exhibited no cytotoxicity. There was significant difference amongst the cytotoxicity of all the extracts (p≤0.05).

### Selectivity index (SI) of *Senna didymobotrya*, *Albizia gummifera* and *Rhamnus staddo*

In *A. gummifera* the greatest SI was observed on the root bark MeOH: DCM extract on the breast cancer cell line (SI = 21.68) and on the colorectal cancer cell line (SI = 8.57). A SI was also recorded on the stem bark MeOH: DCM and aqueous extracts, and root bark MeOH: DCM extract on the prostate cancer cell line (SI = 4.79, 3.44, 3.28), respectively. The stem bark MeOH: DCM and aqueous extracts had a SI of 3.16 and 2.94 on breast and colorectal cancer cell lines respectively. The least selective extracts on the prostate cancer cell line was the root bark MeOH: DCM extract (SI = 0.65), followed by the leaf MeOH: DCM extracts (SI = 1.32) and then the leaf aqueous extract (SI = 1.51). On the breast cancer cell line, the leaf MeOH: DCM extract was the least selective with a SI of 1.50. And finally, on colorectal cancer cell line, the least selective extracts were the root bark aqueous extract (SI = 1.28) and leaf MeOH: DCM extract (SI = 1.07). In *S. didymobotrya*, the root bark MeOH: DCM extract was the most selective with a SI of 6.59 on the colorectal cancer cell line. The least selective extract was the stem bark MeOH: DCM extract on throat cancer cell line with a SI of 1.49. For *R. staddo* root bark MeOH: DCM extract, a high selectivity (SI ≥ 3) was observed on all the cancer cell lines tested (Table 2)

**Table 2:** Selectivity index of Senna didymobotrya, Albizia gummifera and Rhamnus staddo. Key: N/A: Not applicable because the test drug did not inhibit the growth of the cell KEY: MeOH: DCM-Methanol: Dichloromethane (1:1)

| **Plant Sample** | **Part Used** | **Solvent** | **DU 145** | **HCC 1395** | **Hep 2** | **CT 26** |
| --- | --- | --- | --- | --- | --- | --- |
| ***S. didymobotrya*** | Leaf | Aqueous | N/A | N/A | N/A | N/A |
|  | Stem bark | Aqueous | N/A | N/A | N/A | N/A |
|  | Root bark | Aqueous | N/A | N/A | N/A | N/A |
|  | Leaf | MeOH: DCM | 1.52 | 3.09 | 2.52 | N/A |
|  | Stem bark | MeOH: DCM | N/A | 1.70 | 1.49 | N/A |
|  | Root bark | MeOH: DCM | N/A | 1.53 | 1.61 | 6.58 |
| ***A. gummifera*** | Leaf | Aqueous | 1.51 | N/A | N/A | N/A |
|  | Stem bark | Aqueous | 3.44 | 2.94 | 2.66 | 2.33 |
|  | Root bark | Aqueous | 3.28 | 2.33 | 1.49 | 1.28 |
|  | Leaf | MeOH: DCM | 1.32 | 1.59 | 1.72 | 1.07 |
|  | Stem bark | MeOH: DCM | 4.79 | 2.60 | 2.25 | 3.16 |
|  | Root bark | MeOH: DCM | 0.65 | 21.68 | 4.22 | 8.57 |
| ***R. staddo*** | Root bark | MeOH: DCM | 5.15 | 3.03 | 4.81 | 4.54 |

### Assessment of expression of apoptotic (*p53*) and angiogenic (*VEGF*) genes

This study investigated the changes in *p53* and *VEGF* gene expressions in DU145 prostate cancer cells. The relative mRNA expression levels of the genes were determined using a real time PCR. The fold increase or decrease in the expression of the genes was evaluated, relative to the calibrator (Relative Quantification=1). It was observed that all the extracts showed a significant fold increase in the *p53* expression (Table 3). The MeOH: DCM extracts of *S. didymobotrya*, *A. gummifera*, *R. staddo* increased *p53* expression in DU145 by a fold change of 15.990, 16.066 and 15.985 respectively. A downregulation of the *VEGF* gene was noted in cells treated with *S. didymobotrya*, *A. gummifera* and *R. staddo* MeOH: DCM extracts with a fold change of 0.015±0.007, – 0.070±0.015 and 0.045 respectively. (Table 3).

**Table 3:** Fold change in expression of mRNA apoptotic (p53) and angiogenic (VEGF) genes

| Sample | P53 gene expression | VEGF gene expression |
| --- | --- | --- |
| *S. didymobotrya* leaf MeOH: DCM | $15.990 \pm 0.014$ | $0.015 \pm 0.007$ |
| *A. gummifera* stem bark MeOH: DCM | $16.066 \pm 1.965$ | $0.070 \pm 0.015$ |
| *R. staddo* root bark MeOH: DCM | $15.985 \pm 0.015$ | $0.045 \pm 0.017$ |

Key: MeOH: DCM-Methanol dichloromethane (1:1)

### Qualitative Phytochemical Screening

Phytochemical screening demonstrated the presence of different types of phytocompounds including alkaloids, saponins, flavonoids, phenols, glycosides, tannins and terpenoids which could be responsible for the various pharmacological properties. Saponins, flavonoids, glycosides, terpenoids and tannins were found across all the plant extracts. Phenols were also found present in all extracts apart from the root and stem aqueous extracts of *A. gummifera*. Alkaloids were present in *A. gummifera* extracts except in the leaf aqueous extract. Alkaloids were also exhibited in *R. staddo* and *S. didymobotrya* roots methanol dichloromethane extracts (Table 4)

**Table 4:**
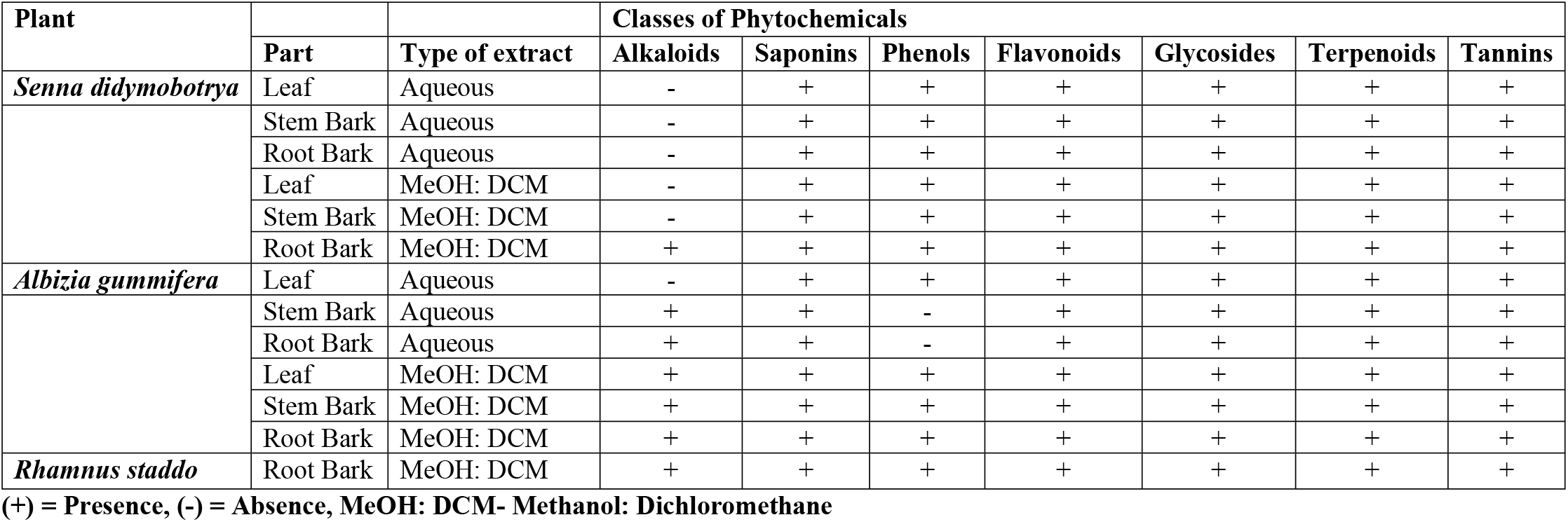
Phytochemical constituents of aqueous and methanol dichloromethane extracts of *S. didymobotrya*, *A. gummifera* and *R. staddo*

| Plant |  |  | Classes of Phytochemicals |  |  |  |  |  |  |
| --- | --- | --- | --- | --- | --- | --- | --- | --- | --- |
|  | Part | Type of extract | Alkaloids | Saponins | Phenols | Flavonoids | Glycosides | Terpenoids | Tannins |
| *Senna didymobotrya* | Leaf | Aqueous | - | + | + | + | + | + | + |
|  | Stem Bark | Aqueous | - | + | + | + | + | + | + |
|  | Root Bark | Aqueous | - | + | + | + | + | + | + |
|  | Leaf | MeOH: DCM | - | + | + | + | + | + | + |
|  | Stem Bark | MeOH: DCM | - | + | + | + | + | + | + |
|  | Root Bark | MeOH: DCM | + | + | + | + | + | + | + |
| *Albizia gummifera* | Leaf | Aqueous | - | + | + | + | + | + | + |
|  | Stem Bark | Aqueous | + | + | - | + | + | + | + |
|  | Root Bark | Aqueous | + | + | - | + | + | + | + |
|  | Leaf | MeOH: DCM | + | + | + | + | + | + | + |
|  | Stem Bark | MeOH: DCM | + | + | + | + | + | + | + |
|  | Root Bark | MeOH: DCM | + | + | + | + | + | + | + |
| *Rhamnus staddo* | Root Bark | MeOH: DCM | + | + | + | + | + | + | + |

(+) = Presence, (-) = Absence, MeOH: DCM- Methanol: Dichloromethane

## Discussion

### Antiproliferative activity of *Senna didymobotrya*, *Albizia gummifera* and *Rhamnus staddo*

Generally, the plant extracts inhibited the proliferation of the cancer cells. The antiproliferative activities of the extracts were categorized based on median inhibitory concentration (IC_50_) [38] Based on the National Cancer Institute (NCI) criterion, *A. gummifera* stem bark MeOH: DCM extracts showed the most potent cytotoxic effect on all the cancer cells tested, with an inhibitory concentration of less than 20μg/ml against all cell lines. The stem bark aqueous extracts also exhibited high cytotoxicity on the prostate cancer cells with an IC_50_ value of 18.29±0.02μg /ml. Notably, the IC_50_ values of *A. gummifera* stem bark MeOH: DCM extract on prostate cancer cells (3.34μg /ml) and *A. gummifera* root bark MeOH: DCM extract on breast cancer cells (2.38μg /ml) were found to be lower than those specified by the NCI, USA for categorization of a pure compound (IC_50_ < 4μg/ml). The *R. staddo* root bark MeOH: DCM extract was classified as highly antiproliferative against all the cell lines with an IC_50_ value less than 20μg /ml, with the highest antiproliferative activity was observed against DU145 cells with an IC_50_ value of 9.36±0.10μg/ml. The MeOH: DCM leaf extracts of *S. didymobotrya* was categorized as highly antiproliferative against colon CT26 cancer cell line with an IC_50_ of 15.13±0.03 μg/ml. The extract reported a moderate antiproliferative activity against DU145 cells (IC_50_ of 65.72± 0.01 μg/ml), HCC 1395 (IC_50_ of 32.32± 0.03 μg/ml) and Hep2 cells (IC_50_ of 39.17±0.29 μg/ml). Doxorubicin was used as the positive control. It showed the highest antiproliferative activity, but it was the most toxic to normal cells.

Several studies have demonstrated the antiproliferative activity of *Albizia* species. The methanolic bark extract of *A. adinocephala* showed antiproliferative activity on breast cancer expressing receptor HER-2+, estrogen dependent breast cancer cells (ER+) and leukemia CCRF-CEM cells with IC_50_ values of 4.8μg/ml, 5.9μg/ml and 1.45μg/ml, respectively [39]. *A. adinocephala* root extract also exhibited excellent cytotoxic effects on leukemia CCRF-CEM cells with an IC_50_ value of 0.98μg/ml [40]. The aqueous and methanolic extracts of *A. amara* exhibited cytotoxic effects to the MCF-7 breast cancer cells with IC_50_ values of 83.16μg/ml and 57.53μg/ml, respectively. A stronger cytotoxicity was found in the ethyl acetate extract with an IC_50_ value of 36.31μg/ml [41]. The cytotoxicity of *A. zygia* hydroethanolic and curcumin extracts has also been reported with IC_50_ values of 3.37μg/ml on (Jurkat) human T-lymphoblast-like leukemia, 6.08μg/ml on (LNCap) prostate cancer cells and 2.56μg/ml on (MCF-7) breast cancer cells [42]. These previous studies demonstrate the potential of Albizia species to inhibit the growth of several cancer cell lines.

A number of species belonging to the Rhamnaceae family have been shown to possess antiproliferative properties on cancer cell lines. *Ziziphus spina Christi* demonstrated antiproliferative potential on MCF-7 breast cancer cells by inducing apoptosis with an IC_50_ value of 20μg/ml [43]. *Rhamnus davunica* exhibited antiproliferative effect on human cancer cells HT-29 (colon carcinoma) and SGC-79 (gastric carcinoma) with IC_50_ values of 24.96μg/ml and 89.53μg/ml, respectively. *Rhamnus nepalensis* is also reported to be cytotoxic to the human oral carcinoma cell line with an IC_50_ value of 35.69μg/ml [44].

A number of *Senna* species have been shown to have antiproliferative activity against several cancer cell lines. The aqueous root bark extract of *Cassia abbreviata* inhibited the growth of Hep2 throat cancer cell line with an IC_50_ value of 1.49μg/ml [45]. *Senna alata* methanolic leaf extract with 22.46μg/ml exhibited growth inhibitory potential on HepG2 liver cancer cell line [46]. The leaf extract of *T. cacao* (Senna spp) exhibited antiproliferative activity against MCF-7 breast cancer cells with an IC_50_ value of 41.4μg/ml. The husk fermented shell of *T. cacao* had IC_50_ values of 71.4μg/ml against HeLa and 68.9μg/ml on HepG2 [47]. On the contrary, *Senna covensii* leaf methanolic was reported to have no antiproliferative activity on HeLa cell line [48].

The anti-proliferation potential of these plants could be attributed to the phytochemicals present. Natural products have been an important source of pharmacologically active molecules that are effective and have a low occurrence of side effects [49]. Phytochemical screening demonstrated the presence of different types of phytocompounds including alkaloids, saponins, flavonoids, phenols, glycosides tannins and terpenoids. Phytochemical studies of different species belonging to the *Albizia* genus has revealed different classes of secondary metabolites such as saponins, terpenes, alkaloids and flavonoids [10]. In this study, all the methanol dichloromethane extracts of *A. gummifera* contained alkaloids, saponins, terpenoids, flavonoids and phenols, results that correlate with the findings done by Nigussie *et al* [50]. Phenols were absent in the root and stem aqueous extracts of *A. gummifera*.

Saponins, flavonoids, sterols, alkaloids, phenol, tannins, terpenoids and glycosides were found in leaf, stem bark and root bark aqueous and MeOH: DCM extracts of *S. didymobotrya*. According to reports on pharmacological activity of Senna species, it has been suggested that the genus is a potential source of enormous bioactive compounds. Phytochemicals such as tannins, alkaloids, saponins, steroids, flavonoids, and terpenoids are constituents of *S. didymobotrya* [51].

Similarly, the methanol dichloromethane roots extracts of *S. didymobotrya* had all these phytochemicals. This correlated with a study that reported the presence of tannins, alkaloids, saponins, steroids, flavonoids, and terpenoids in the roots extracts of *S. didymobotrya*. [52]. Another study also reported the presence of saponins, terpenoids and flavonoids in the stem and root extracts of *S. didymobotrya* [53]. The leaf extract also revealed the presence of tannins, saponins, flavonoids, glycosides, sterols and phenols [53]. The leaf however did not have alkaloids. This was in line with a study conducted by Ngule *et al* [54]. Likewise, all these phytochemicals were found in *S. didymobotrya* extracts in the present study.

Saponins, flavonoids, sterols, alkaloids, terpenoids and glycosides were found present in the *R. staddo* root methanol dichloromethane extract. The root extract of *R. staddo* has similarly been reported to be rich in saponins, flavonoids, sterols, alkaloids, terpenoids and glycosides [55]. Additionally, *Ziziphus lotus* (*Rhamnacea*) was observed to contain similar phytochemical profiles including alkaloids, saponins, terpenoids, flavonoids and coumarins. However, the extract did not contain anthraquinones [56].

A number of studies have been conducted to prove the protective effect of flavonoids against cancer. It has been linked to reduced incidence of estrogen-related cancers [57,58]. A correlation has been observed between the flavonoids in *R. davunica* with its antiproliferative activities against colon and gastric cancer cells [59]. *S. didymobotrya* was found to have flavonoids [44]. These are secondary metabolites that are potent water-soluble antioxidants and free radical scavengers that prevent oxidative cell damage and have strong anticancer activity [60]. Increased consumption of isoflavones is directly proportional to decreased risk of cancer and vascular diseases [61], with a 50% reduction in the risk of being susceptible to stomach and lung cancer [62]. *In vitro*, isoflavonoids inhibited the proliferation of breast cancer cells in a concentration dependent manner [63]. Flavonoids have also been pointed out as enzyme inhibitors and ligands of receptors involved in signal transductions [64,65].

Terpenoids have been demonstrated to possess antitumor properties, which has attracted more attention and these compounds have been screened in multiple cultured cancer cells. Several terpenoids have been isolated from plants and have been proved to have antiproliferative potential [66]. Terpenoids isolated from *Eurphobia kansui* exerted an antiproliferative effect on MDA-MB-435 breast cancer cells and Colo205 colorectal cancer cells [67]. The terpenoids extracted from *Ferulago macrocarpa* fruits exerted cytotoxic effects on MCF-7 breast cancer cells and HT-27 colon cancer cells [68].

Alkaloids have been reported to be important active components in medicinal plants with significant biological activities. Some of these compounds have been developed successfully into chemotherapeutic drugs such as vinblastine, camptothecin and topoimerase 1 [69]. *Albizia amara’s* seeds methanolic extract was found to contain macrocycic pithecolobine alkaloids which rendered its high cytotoxic potential towards human breast cancer, colon cancer, lung cancer and melanoma cell lines [70]. Noscapine, an alkaloid isolated from the opium flower *Papaver somniferum L*. induced G2/M arrest in colorectal cancer [71]. The activity of the root extract of *S. didymobotrya* could therefore be attributed to the alkaloids present. Saponins are natural glycosides, possessing a wide range of pharmacological properties including cytotoxicity, antitumor, immunomodulatory, antifungal and antiparasitic [72]. Oleanane-type triterpenoid saponins isolated from the roots of *A. gummifera* after a bio-assay guided fractionation showed cytotoxicity against the A2780 human ovarian cancer cell lines [12]. A triterpene saponin enriched fraction from *Bupleurum kaoi* inhibited the growth of lung cancer A549 cells through the apoptotic pathways with an enhancement in Fas ligand [73]. Cardiac glycosides tend to exert potent antineoplastic effects by increasing the immunogenicity of dying cancer cells [74].

The selective inhibitory activity of the extracts was determined and expressed as selectivity index (SI). The SI values demonstrated the differential activity of the extracts on normal cells compared to cancerous cell lines. A high SI value depicts high selectivity. Medicinal plants with SI values of 2 or greater than 2 are considered to be highly selective. Selectivity index of less than 2 indicates less selectivity [75].

*A. gummifera* showed selective toxicity to the cancer cells while sparing the Vero (normal) cells. However, the leaf MeOH: DCM extract exhibited non-selective inhibitory activity across all the cell lines tested with a selectivity index less than 2. This is similar to the findings in a study where the leaf of *A. adianthifolia* displayed toxicity to normal AML12 hepatocytes [76]. The root aqueous extracts were also found to be toxic to the normal cells with SI of 1.49 and 1.28 on the throat and colorectal cancer cell lines, respectively. The lowest selectivity was exhibited by the root bark MeOH: DCM extract with a SI of 0.65 on the prostate cancer cells. A study on *A. zygia* root exhibited toxicity on breast cancer cells with SI of 0.72 on the aqueous extract, 0.75 on the hydroethanolic extract and 1.86 on the curcumin extract. This was however different in *A. gummifera* root MeOH: DCM extract that had the highest selectivity on the breast cancer cells with a SI of 21.68. Nevertheless, the toxicity of these extracts towards cancer cells makes them potential candidates as anticancer therapeutic agents.

*R. staddo* MeOH: DCM root extract also demonstrated great selectivity to the cancer cells while sparing the normal cells (SI≥3). The highest selectivity was observed on the prostate cancer cells with a SI of 5.15. This is parallel to the findings of Chen *et al*., where *R. davunica* showed no toxicity on the normal human hepatic cells (L-O2) [55].

*S. didymobotrya* extracts moderately inhibited the growth of cancer cells and were non-toxic to normal cells. The leaf MeOH: DCM extracts showed selectivity on both breast and throat cancer cells with SI of 3.09 and 2.52, respectively. The root bark MeOH: DCM extract had the best activity which was expressed on colorectal cancer cells an activity that was not exhibited by the other *Senna* extracts. It also expressed high selectivity to the colorectal cancer cells (SI= 6.58). This potentiates its use in colorectal cancer management because it showed strong cytotoxicity to the cancer cells while being non-toxic to the normal cells. The finding of the preset study concurs with the results showing that *Cassia abbreviata* selectively inhibited Hep 2 cancer cell lines while sparing the normal cell line [77]. This study reports for the first time the anti-proliferative potential of *S. didymobotrya*, *A gummifera* and *R staddo* on breast, prostrate, colorectal and throat cancer cells.

The MeOH: DCM extracts of *S. didymobotrya*, *A. gummifera*, *R. staddo* increased *p53*expression in DU145 by a fold change of 15.99, 16.07 and 15.98 respectively. A downregulation of the *VEGF* gene was also noted in the cells treated with *S. didymobotrya* and *A. gummifera* and *R. staddo* extracts with a fold change of 0.015, 0.070 and 0.045, respectively. The medicinal bioactivity of these plants could be related to the identified phytochemicals. A phytomolecule can suppress malignant transformation of an initiated pre-neoplastic cell or can block the metabolic conversion of the pro-carcinogen. These mechanisms of action could also be attributed to the vast range of phytochemicals present in the extracts. Several studies have shown the impact of these isolated phytocompounds to the various mechanisms of plants in inhibition of growth of cancer cells. Polyphyllin D is a steroidal saponin isolated from *Paridis rhizome* and has been shown to inhibit cell proliferation of cancer cells by inhibiting the expression of *VEGF* genes and by upregulating p21 [78]. Ellagic acid of pomegranate induces apoptosis in prostate and breast cancer cells and inhibits metastasis process of various cancer types [79]. Asiatic acid, a pentacyclic triterpene isolated from *Centella asiatica* and oridonin, a diterpene, both inhibited the growth of HepG2 liver cancer cells by increasing the expression of p53 gene [80]. Apigenin, a flavone found in parsley, cereley and chamomile targets the p53 pathway and induces apoptosis in the lung adenocarcima cells. It induces capsase dependent extrinsic apoptosis in human epidermal growth factor receptor 2 (HER-2) over expressing BT-474 breast cancer cells [81]. The proapoptotic effects of Eupatilin, an active flavone derived from *A. asiatica* on human gastric cancer cells has been reported. It was found that the treatment of the human gastric cancer cells with eupatilin resulted in elevated expression of the p53 gene [82].

Flavonoids also act as anti-angiogenic compounds by downregulating the expression of *VEGF* gene. Crocetin, a caretonoid in *Crocus sativus* and *Gardenia jasminoides* suppresses the production of the *VEGF* gene in the head and neck carcinoma cells [83]. The diterpene cafestrol from *Coffea arabita* and phenols such as carnosic acid and carnosol have been shown to inhibit angiogenesis hence preventing the process of carcinogenesis [84]. Curcumin, a polyphenol of *curcuma longa* inhibits the growth of human glioblastoma cells by upregulating the expression of p53 gene [85]. Triterpenoids that enhance cytotoxicity to cancer cells causes an increase in the level of reactive oxygen species leading to increased apoptosis [86]. Studies have shown that a combination of phytochemicals can be more effective against cancer than individual components [86]. The probable mechanism of action of *S. didymobotrya*, *A. gummifera*, *R. staddo* could be attributed to the phytochemicals present.

## Conclusion

The plant extracts could be potential candidates for development of drugs for the management of breast, prostrate, colorectal and throat cancer. Induction of apoptosis and anti-angiogenesis could be proposed as their probable mechanisms of action. The growth inhibitory potential of the plant extracts on the cancer cells and the probable mechanism of action could be attributed to the presence of pharmacologically important phytochemicals. This study confirms that amidst the many traditional and pharmacological uses of these plants, they could also be used in the fight against cancer menace.

## Conflict of Interest

The authors declare no conflicts of interests

## Ethical Consideration

There were no humans involved in this study. The animals and cell lines used were handled with a lot of care and professionalism and all protocols followed to the letter. All the safety standards in the place of study were observed and all measures considered to make sure that standard operating procedures were carried out maximally. Ethical approval was sought from Kenyatta University Graduate School Committee, Kenya Medical Research Institute (KEMRI), CTMDR Centre Scientific Committee (CSC) and Scientific and Ethics Review Unit (SERU) approval number KEMRI/SERU/CTMDR/O55/3535 before conducting the study

## Acknowledgements

We are grateful to Kenya Medical Research Institute, Centre for Traditional Medicine and Drug Research fraternity for their unending support, guidance and co-operation in actualizing this work. We acknowledge the KEMRI’s Internal Research Grant (IRG) for funding Dr. Peter Mwitari through SSC 2902 who supplied us with laboratory consumables and the cancer cell lines. We are also thankful to the Pyrethrum project funding, SSC 2969 that facilitated the purchase of the q-PCR and CO_2_ incubator, equipment’s that aided in the success of this work.

## References

1. WHO. World cancer report. The International Agency for Research on Cancer. Lyon, London. 2014

2. Korir A, Okerosi N, Ronoh V, Mutuma G, Parkin M. Incidence of cancer in N airobi, K enya (2004–2008). International journal of cancer. 2015 Nov 1;137(9):2053–9.

3. Joy P, Thomas J, Mathew S, Skaria B. Medicinal plants in Tropical Horticulture, vol.2, page .449–632, 2001 NayaProkash,Calcutta, India

4. Aruna MS, Prabha MS, Priya NS, Nadendla R. Catharanthus Roseus: ornamental plant is now medicinal boutique. Journal of Drug Delivery and therapeutics. 2015 Feb 10;5(3):1–4.

5. Tabuti JR. Senna didymobotrya (Fresen). HS Irwin barneby. Schmeltzer, GH Gurib. Fakim, A (editors) Prota. 2007;11.

6. Ochwang’I DO, Kimwele CN, Oduma JA, Gathumbi PK, Kiama SG. Phytochemical Screening of Medicinal Plants of the Kakamega Country, Kenya Commonly Used against Cancer. Medicinal and Aromatic Plants (Los Angel). 2016;5(277):2167–0412.

7. Korir RK, Mutai C, Kiiyukia C, Bii C. Antimicrobial activity and safety of two medicinal plants traditionally used in Bomet District of Kenya. Research journal of Medicinal plants. 2012;6(5):370–82.

8. Nagappan R. Evaluation of aqueous and ethanol extract of bioactive medicinal plant, Cassia didymobotrya (Fresenius) Irwin & Barneby against immature stages of filarial vector, Culex quinquefasciatus Say (Diptera: Culicidae). Asian pacific Journal of tropical Biomedicine. 2012 Sep 1;2(9):707–11.

9. Schmidt LH, Mwaura L. Albizia gummifera (JF Gmel.) CA Sm. Seed Leaflet. 2010(141).

10. Kokila K, Priyadharshini SD, Sujatha V. Phytopharmacological properties of Albizia species: a review. International Journal of Pharmacy and Pharmaceutical Sciences. 2013;5(3):70–3.

11. Tadesse, E. In Vitro and in Vivo Evaluation of Anthelmintic Activities of Crude Extracts of Selected Medicinal Plants against Haemonchus Contortus. Medicinal and Aromatic Plants 2005

12. Cao S, Norris A, Miller JS, Ratovoson F, Razafitsalama J, Andriantsiferana R, Rasamison VE, TenDyke K, Suh T, Kingston DG. Cytotoxic Triterpenoid Saponins of Albizia gummifera from the Madagascar Rain Forest. Journal of natural products. 2007 Mar 23;70(3):361–6.

13. Veeresh B, Babu SV Patil AA, Warke YB. Lauric acid and myristic acid prevent testosterone induced prostatic hyperplasia in rats. European journal of pharmacology. 2010 Jan 25;626(2–3):262–5.

14. Farmani F, Moein M, Amanzadeh A, Kandelous HM, Ehsanpour Z, Salimi M. Antiproliferative evaluation and apoptosis induction in MCF-7 cells by *Ziziphus spina christi* leaf extracts. Asian Pacific journal of cancer prevention. 2016;17(1):315–21.

15. Duncan C. M. An ethnobotanical study of medicinal plants used by the Masaai people of Losho, Kenya. International Journal of Production Research, 2016 6(02), 68.

16. Onyango CA, Gakuya LW, Mathooko FM, Maina JM, Nyaberi M, Makobe M, Mwaura F. Phytochemical studies on herbal plants commonly used for processing and preserving meat and milk. Journal of Applied Biosciences. 2014;73(1):5942–8.

17. Muregi FW, Ishih A, Miyase T, Suzuki T, Kino H, Amano T, Mkoji GM, Terada M. Antimalarial activity of methanolic extracts from plants used in Kenyan ethnomedicine and their interactions with chloroquine (CQ) against a CQ-tolerant rodent parasite, in mice. Journal of Ethnopharmacology. 2007 Apr 20;111(1):190–5.

18. Vijayalakshmi R, Ravindhran R. HPTLC method for quantitative determination of gallic acid in ethanolic root extract of Diospyrus ferrea (Willd.) Bakh and Aerva lanata (L.) Juss. Ex Schultes—a potent indian medicinal plants. Asian Journal of Pharmaceutical and Clinical Research. 2012;5(4):170–4.

19. Doss A. Preliminary phytochemical screening of some Indian medicinal plants. Ancient science of life, 29(2), 12. 2009

20. Pandey P, Mehta R, Upadhyay R. Physico-chemical and preliminary phytochemical screening of Psoralea corylifolia. Archieve Applied Science Research. 2013;5(2):261–5.

21. Stoff-Khalili MA, Dall P, Curiel DT. Gene therapy for carcinoma of the breast. Cancer gene therapy. Journal of Ethnopharmacology 2006 Jul;13(7):633.

22. Aderonke ST, Babatunde JA, Adesola OT, Okereke OU, Innocent C, Elisha MO, Abolaji OL, Abiola MO. Evaluation of retinoblastoma (Rb) and protein-53 (p53) gene expression levels in breast cancer cell lines (MCF-7) induced with some selected cytotoxic plants. Journal of Pharmacognosy and Phytotherapy. 2013 Jul 31;5(7):120–6.

23. Bieging KT, Mello SS, Attardi LD. Unravelling mechanisms of p53-mediated tumour suppression. Nature Reviews Cancer. 2014 May;14(5):359.

24. Zhang HT, Luo H, Wu J, Lan LB, Fan DH, Zhu KD, Chen XY, Wen M, Liu HM. Galangin induces apoptosis of hepatocellular carcinoma cells via the mitochondrial pathway. World journal of gastroenterology: WJG. 2010 Jul 21;16(27):3377.

25. Sagar SM, Yance D, Wong RK. Natural health products that inhibit angiogenesis: a potential source for investigational new agents to treat cancer—Part 1. Current Oncology. 2006 Feb;13(1):14.

26. Izawa JI, Dinney CP. The role of angiogenesis in prostate and other urologic cancers: a review. Canadian Medical Association Journal. 2001 Mar 6;164(5):662–70.

27. Ferrara, N. Role of vascular endothelial growth factor in regulation of physiological angiogenesis. American Journal of Physiology. 2001 280(76):1358–1366.

28. Leyon, PV. and Kuttan, G. Effect of Tinospora cordifolia on the cytokine profile of angiogenesis-induced animals. International Immunopharmacology. 2004 4(7):1569–1575.

29. Thippeswamy, G., Sheela, ML. and Salimath, BO. Octacosanol isolated from Tinospora cordifolia down regulates VEGF gene expression by inhibiting nuclear translocation of NF-kB and its DNA binding activity. European Journal, 2008 107:308–314.

30. Christina, AJ., Joseph, DG., Packialakshmi, M., Kothai R. and Robert SJ. Evaluation of the effect of Withania somnifera root extracts on cell cycle and angiogenesis. Ethnopharmacoly, 2006 105(3):336–41.

31. Xio, D. and Singh S.V., 2008. Z-guggulsterone a constituent of Ayurvedic medicinal plant *Commiphora mukul* inhibits angiogenesis *in vitro* and *in vivo*. Molecular Cancer Therapeutics, 2008 7(2):171–180.

32. Izawa, JI. and Dinney, CP. The role of angiogenesis in prostate and other urologic cancers: a review. Canadian Medical Association Journal, 2001 164(5):662–70.

33. Donkor, PO., Donnapee, S., Li, J., Yang, X., Ge, AH., Donkor, PO, Gao XM, Chang YX. Cuscuta chinensis Lam.: A systematic review on ethnopharmacology, phytochemistry and pharmacology of an important traditional herbal medicine. Journal of ethnopharmacology, 2101 9(05):98.

34. Mwitari PG, Ayeka PA, Ondicho J, Matu EN, Bii CC. Antimicrobial activity and probable mechanisms of action of medicinal plants of Kenya: Withania somnifera, Warbugia ugandensis, Prunus africana and Plectrunthus barbatus. PloS one. 2013 Jun 13;8(6):e65619.

35. Swamy T, Ngule MC, Jackie K, Edwin A, Ngule ME. Evaluation of in vitro antibacterial activity in Senna didymobotrya roots methanolic-aqua extract and the selected fractions against selected pathogenic microorganisms. International Journal of Current Microbiology and Applied Sciences 2014;3(5):362–76.

36. Nemati F, Dehpouri AA, Eslami B, Mahdavi V, Mirzanejad S. Cytotoxic properties of some medicinal plant extracts from Mazandaran, Iran. Iranian Red Crescent Medical Journal. 2013 Nov;15(11).

37. Yuan JS, Reed A, Chen F, Stewart CN. Statistical analysis of real-time PCR data. BMC bioinformatics. 2006 Dec;7(1):85.

38. Alshatwi AA, Hasan TN, Shafi G, Syed NA, Al-Assaf AH, Alamri MS, Al-Khalifa AS. Validation of the antiproliferative effects of organic extracts from the green husk of Juglans regia L. on PC-3 human prostate cancer cells by assessment of apoptosis-related genes. Evidence-Based Complementary and Alternative Medicine. 2012;2012.

39. Boik J. Natural compounds in cancer therapy. 2001

40. Agudo DA, Paredes M, Shantall N, Espinosa Rivas AF, Guerra Torres CP, Prashad Gupta M. Evaluation of the antiproliferative activity of methanolic extracts of plants from the Leguminosae family. Revista Cubana de Plantas Medicinales. 2017 Mar 27;21(3):272–83.

41. Kuete V, Tchinda CF, Mambe FT, Beng VP, Efferth T. Cytotoxicity of methanol extracts of 10 Cameroonian medicinal plants towards multi-factorial drug-resistant cancer cell lines. BMC complementary and alternative medicine. 2016 Dec;16(1):267.

42. Badisa RB, Darling-Reed SF, Joseph P, Cooperwood JS, Latinwo LM, Goodman CB. Selective cytotoxic activities of two novel synthetic drugs on human breast carcinoma MCF-7 cells. Anticancer research. 2009 Aug 1;29(8):2993–6.

43. Appiah-Opong RE, Asante IK, Safo DO, Tuffour IS, Ofori-attah EB, Uto TA, Nyarko AK. Cytotoxic effects of Albizia zygia (DC) JF Macbr, a Ghanaian medicinal plant, against human t-lymphoblast-like leukemia, prostate and breast cancer cell lines. International Journal of Pharmacy and Pharmaceutical Sciences. 2016;8(5):392–6.

44. Farmani F, Moein M, Amanzadeh A, Kandelous HM, Ehsanpour Z, Salimi M. Antiproliferative evaluation and apoptosis induction in MCF-7 cells by Ziziphus spina christi leaf extracts. Asian Pacific Journal of Cancer Prevention. 2016;17(1):315–21.

45. Mai LP, Guéritte F, Dumontet V, Tri MV, Hill B, Thoison O, Guénard D, Sévenet T. Cytotoxicity of Rhamnosylanthraquinones and Rhamnosylanthrones from Rhamnus nepalensis. Journal of natural products. 2001 Sep 28;64(9):1162–8.

46. Njagi, S.M., Lagat, R.C., Mawia, A.M., Arika, W.M., Wambua, F.K., Ouko, R.O., Ogola, P.E., Kamau, J.K., Nthiga, P.M., Musila, M.N. and Mwitari, P.G. In Vitro Antiproliferative Activity of Aqueous Root Bark Extract of Cassia abbreviata (Holmes) Brenan. Journal of cancer science and therapy, 2016

47. Manosroi A, Akazawa H, Kitdamrongtham W, Akihisa T, Manosroi W, Manosroi J. Potent Antiproliferative Effect on Liver Cancer of Medicinal Plants Selected from the Thai/Lanna Medicinal Plant Recipe Database “MANOSROI III”. Evidence-Based Complementary and Alternative Medicine. 2015

48. Baharum Z, Akim AM, Taufiq-Yap YH, Hamid RA, Kasran R. In vitro antioxidant and antiproliferative activities of methanolic plant part extracts of Theobroma cacao. Molecules. 2014 Nov 10;19(11):18317–31.

49. Estrada M, Velázquez-Contreras C, Garibay-Escobar A, Sierras-Canchola D, Lapizco-Vázquez R, Ortiz-Sandoval C, Burgos-Hernández A, Robles-Zepeda RE. In vitro antioxidant and antiproliferative activities of plants of the ethnopharmacopeia from northwest of Mexico. BMC complementary and alternative medicine. 2013 Dec;13(1):12.

50. Newman DJ, Cragg GM. Natural products as sources of new drugs over the 30 years from 1981 to 2010. Journal of natural products. 2012 Feb 8;75(3):311–35

51. Nigussie D, Tasew G, Makonnen E, Debella A, Hurrisa B. In-vitro investigation of fractionated extracts of Albizia gummifera seed against Leishmania donovani amastigote stage. Journal of Clinical and Cellular Immunology. 2015;6(373):2.

52. Nyamwamu, L.B., Ngeiywa, M., Mulaa, M. and Lelo, A.E. Phytochemical constituents of senna didymobotrya fresen irwin roots used as a traditional medicinal plant in kenya., Plant Product Research Journal, 13, 2015 pp.35–43.

53. Kitonde CK, Fidahusein D, Lukhoba CW, Jumba MM. Antimicrobial activity and phytochemical screening of Senna didymobotry used to treat bacterial and fungal infections in Kenya. International Journal of Education and Research. 2014;2(1):1–2.

54. Ngule CM. Phytochemical and bioactivity evaluation of Senna didymobotrya fresen irwin used by the Nandi community in Kenya. International Journal of Bioassays. 2013 Jun 27;2(7):1037–43.

55. Onyango CA, Gakuya LW, Mathooko FM, Maina JM, Nyaberi M, Makobe M, Mwaura F. Phytochemical studies on herbal plants commonly used for processing and preserving meat and milk. Journal of Applied Biosciences. 2014;73(1):5942–8.

56. Naili MB, Alghazeer RO, Saleh NA, Al-Najjar AY. Evaluation of antibacterial and antioxidant activities of Artemisia campestris (Astraceae) and Ziziphus lotus (Rhamnacea). Arabian Journal of Chemistry. 2010 Apr 1;3(2):79–84.

57. Pietinen P, Stumpf K, Männistö S, Kataja V, Uusitupa M, Adlercreutz H. Serum enterolactone and risk of breast cancer: a case-control study in eastern Finland. Cancer Epidemiology and Prevention Biomarkers. 2001 Apr 1;10(4):339–44.

58. Dai Q, Franke AA, Jin F, Shu XO, Hebert JR, Custer LJ, Cheng J, Gao YT, Zheng W. Urinary excretion of phytoestrogens and risk of breast cancer among Chinese women in Shanghai. Cancer Epidemiology and Prevention Biomarkers. 2002 Sep 1;11(9):815–21.

59. Chen G, Li X, Saleri F, Guo M. Analysis of flavonoids in rhamnus davurica and its antiproliferative activities. Molecules. 2016 Sep 23;21(10):1275.

60. Yadar

61. Arai Y, Watanabe S, Kimira M, Shimoi K, Mochizuki R, Kinae N. Dietary intakes of flavonols, flavones and isoflavones by Japanese women and the inverse correlation between quercetin intake and plasma LDL cholesterol concentration. The Journal of nutrition. 2000 Sep 1;130(9):2243–50.

62. Neuhouser ML. Dietary flavonoids and cancer risk: evidence from human population studies. Nutrition and cancer. 2004 Sep 1;50(1):1–7.

63. Mueller SO, Simon S, Chae K, Metzler M, Korach KS. Phytoestrogens and their human metabolites show distinct agonistic and antagonistic properties on estrogen receptor α (ERα) and ERβ in human cells. Toxicological Sciences. 2004 Apr 14;80(1):14–25.

64. Williams RJ, Spencer JP, Rice-Evans C. Flavonoids: antioxidants or signalling molecules?. Free radical biology and medicine. 2004 Apr 1;36(7):838–49.

65. Brown AK, Papaemmanouil A, Bhowruth V, Bhatt A, Dover LG, Besra GS. Flavonoid inhibitors as novel antimycobacterial agents targeting Rv0636, a putative dehydratase enzyme involved in Mycobacterium tuberculosis fatty acid synthase II. Microbiology. 2007 Oct 1;153(10):3314–22.

66. Huang J, Wu L, Tashiro SI, Onodera S, Ikejima T. Reactive oxygen species mediate oridonin-induced HepG2 apoptosis through p53, MAPK, and mitochondrial signaling pathways. Journal of pharmacological sciences. 2008;107(4):370–9.

67. Jin-Jun HO, Yao SH, Zhou YA, Lin FA, Lu-Ying CA, Shuai YA, Hua-Li LO, Wan-Ying WU, De-An GU. Anti-proliferation activity of terpenoids isolated from Euphorbia kansui in human cancer cells and their structure-activity relationship. Chinese journal of natural medicines. 2017 Oct 1;15(10):766–74.

68. Sajjadi SE, Jamali M, Shokoohinia Y, Abdi G, Shahbazi B, Fattahi A. Antiproliferative evaluation of terpenoids and terpenoid coumarins from Ferulago macrocarpa (Fenzl) Boiss. fruits. Pharmacognosy research. 2015 Oct;7(4):322.

69. Lee MR. The history of Ephedra (ma-huang). JR Coll Physicians of Edinburgh 2011 Mar;41(1):78–84

70. Lu JJ, Bao JL, Chen XP, Huang M, Wang YT. Alkaloids isolated from natural herbs as the anticancer agents. Evidence-Based Complementary and Alternative Medicine. 2012;2012.

71. Kumar,2012

72. DeBono A, Capuano B, Scammells PJ. Progress toward the development of noscapine and derivatives as anticancer agents. Journal of medicinal chemistry. 2015 Apr 16;58(15):5699–727.

73. Wong FC, Woo CC, Hsu A, Tan BK. The anti-cancer activities of Vernonia amygdalina extract in human breast cancer cell lines are mediated through caspase-dependent and p53-independent pathways. PLoS One. 2013 Oct 24;8(10):e78021.

74. Menger L, Vacchelli E, Kepp O, Eggermont A, Tartour E, Zitvogel L, Kroemer G, Galluzzi L. Trial watch: cardiac glycosides and cancer therapy. Oncoimmunology. 2013 Feb 1;2(2):e23082.

75. Lee MS, Chan JY, Kong SK, Yu B, Eng-Choon VO, Nai-Ching HW, Mak Chung-Wai T, Fung KP. Effects of polyphyllin D, a steroidal saponin in Paris polyphylla, in growth inhibition of human breast cancer cells and in xenograft. Cancer biology and therapy. 2005 Nov 1;4(11):1248–54.

76. Ma DD, Lu HX, Xu LS, Xiao W. Polyphyllin D exerts potent anti-tumour effects on Lewis cancer cells under hypoxic conditions. Journal of International Medical Research. 2009 May;37(3):631–40.

77. Singh, S., Sharma, B., Kanwar, S.S, Kumar, A. Lead phytochemicals for anticancer drug development. Frontiers in plant science, 7, 2016 p.1667.

78. Lee YS, Jin DQ, Kwon EJ, Park SH, Lee ES, Jeong TC, Nam DH, Huh K, Kim JA. Asiatic acid, a triterpene, induces apoptosis through intracellular Ca2+ release and enhanced expression of p53 in HepG2 human hepatoma cells. Cancer letters. 2002 Dec 1;186(1):83–91.

79. Huang J, Wu L, Tashiro SI, Onodera S, Ikejima T. Reactive oxygen species mediate oridonin-induced HepG2 apoptosis through p53, MAPK, and mitochondrial signaling pathways. Journal of pharmacological sciences. 2008;107(4):370–9.

80. Seo HS, Ku JM, Choi HS, Choi YK, Woo JK, Jo JK, Nam KW, Park N, Jang BH, Shin YC, Ko SG. Induction of caspase-dependent extrinsic apoptosis by apigenin through inhibition of signal transducer and activator of transcription 3 (STAT3) signaling in HER2-overexpressing BT-474 breast cancer cells. Bioscience reports. 2015 Oct 23: BSR20150165.

81. Choi EJ, Oh HM, Na BR, Ramesh TP, Lee HJ, Choi CS, Choi SC, Oh TY, Choi SJ, Chae JR, Kim SW. Eupatilin protects gastric epithelial cells from oxidative damage and down-regulates genes responsible for the cellular oxidative stress. Pharmaceutical research. 2008 Jun 1;25(6):1355–64.

82. Luthra PM, Lal N. Prospective of curcumin, a pleiotropic signalling molecule from Curcuma longa in the treatment of Glioblastoma. European journal of medicinal chemistry. 2016 Feb 15; 109:23–35.

83. Moeenfard M, Cortez A, Machado V, Costa R, Luís C, Coelho P, Soares R, Alves A, Borges N, Santos A. Anti-Angiogenic Properties of Cafestol and Kahweol Palmitate Diterpene Esters. Journal of cellular biochemistry. 2016 Dec;117(12):2748–56.

84. Vallianou, N. G., Evangelopoulos, A., Schizas, N., Kazazis, C. Potential anticancer properties and mechanisms of action of curcumin. Anticancer Research 2015 .35, 645–651

85. Randhawa, M.A. and Alghamdi, M.S., 2011. Anticancer activity of Nigella sativa (black seed)—a review. The American journal of Chinese medicine, 39(06), pp.1075–1091.

86. Liu RH. Potential synergy of phytochemicals in cancer prevention: mechanism of action. The Journal of nutrition. 2004 Dec 1;134(12):3479S–85S.

